# Electrophysiological investigation of reward anticipation and outcome evaluation during slot machine play

**DOI:** 10.1101/2020.10.08.330654

**Authors:** SL Fryer, BJ Roach, CB Holroyd, MP Paulus, K Sargent, A Boos, JM Ford, DH Mathalon

**Affiliations:** Psychiatry, University of California, San Francisco, San Francisco, CA, USA; San Francisco VA Health Care System, San Francisco, CA, USA; Department of Experimental Psychology, Ghent University, Belgium; Laureate Institute for Brain Research, Tulsa, Oklahoma, USA

**Keywords:** feedback related negativity (FRN), delta oscillations, frontal medial theta oscillations, near miss, frustrative nonreward

## Abstract

Slot machines are a popular form of gambling, offering a tractable way to experimentally model reward processes. This study used a 3-reel slot paradigm to assess psychologically distinct phases of reward processing, reflecting anticipation, and early and late-stage outcome processing. EEG measures of winning, nearly missing (a losing outcome revealed at the final, third reel), and “totally” missing (a losing outcome revealed earlier, at the second reel) were collected from healthy adults (n=54). Condition effects were evaluated in: i) event-related potential (ERP) components reflecting anticipatory attention (stimulus preceding negativity, SPN) and outcome processing (reward positivity, RewP and late-positive potential, LPP) and ii) total power and phase synchrony of theta and delta band oscillations. Behaviorally, trial initiation was fastest after a near miss outcome and slowest after a winning outcome. As expected, a significant SPN was observed for possible wins (AA) vs. total misses (AB), consistent with reward anticipation. In addition, significantly larger win (AAA) vs. near miss (AAB) amplitudes were observed for the RewP and LPP to wins and LPP to near misses (vs. total misses) reflecting early and late-stage outcome processing effects. There was an effect of reel position on the RewP, with larger effects in the final reel (AAA-AAB) relative to the 2^nd^-reel locked difference waves (AA-AB). Across all outcomes, near misses elicited the largest and most phase-synchronized theta responses, while wins elicited larger and more phase-synchronized delta responses than total misses, with near misses not differing from wins or total misses. Phase locking measures contrasting win vs. near miss delta and theta synchronization, within time windows corresponding to ERP measurements, covaried with RewP, but not SPN or LPP, amplitude. Lastly, EEG measures showed differential relationships with age and self-reported consummatory pleasure. In the context of slot machine play, where reward anticipation and attainment place minimal demands on effort and skill, ERP and time-frequency methods capture distinct neurophysiological signatures of reward anticipation and outcome processing.

## Introduction

Reward processing is a broad term that encompasses more specific sub-components that are increasingly appreciated on the basis of distinct underlying neurophysiological signatures. One major conceptual distinction parses reward-related functions into anticipatory processes related to reward responsiveness, motivation, and goal-oriented behaviors that have been referred to as “wanting,” relative to consummatory processes related to reward attainment that have been referred to as “liking” ^1^. This distinction is reflected in dissociable neurocircuitry, with “wanting” responses being more associated with distributed neuroanatomical circuitry mediated by dopaminergic neurotransmission and “liking” responses being more associated with GABA, opioid, and endocannabinoid signaling ^2^.

Developing a more complete understanding of how reward-related neural measures relate to each other and underlie psychologically distinct sub-stages of reward processing is important for characterizing reward responsivity in health and disease. Electroencephalography (EEG) can be used to parse in vivo human brain activity into constituent psychological processes with high temporal precision across phases of reward processing. However, to date, many studies of reward-related EEG response have focused more narrowly on a single measure or time point ^3^. We therefore combined traditional time-domain event-related potential (ERP) measures and time-frequency neuro-oscillatory measures of reward anticipation and early and late stages of reward outcome processing.

Electrophysiological measures of reward processing include the reward positivity (RewP) an ERP component that is larger to wins than losses ^4^. The RewP has a frontocentral topography, peaking ∼250-300 ms after external feedback indicating reward outcome ^5-8^, with simultaneously acquired fMRI demonstrating that feedback-locked RewP amplitudes covary with BOLD activations in ventral striatum, anterior cingulate cortex (ACC), and medial prefrontal cortex ^9,10^. The RewP had often been referred to as the feedback-related negativity (FRN), an evaluative component first elicited by feedback-locked analysis of errors in a time estimation task ^11^ which was subsequently shown to be larger to losses than wins in rewarded paradigms ^4^. Because much of the ERP variability in this waveform appeared to derive from win events, with the loss-related negativity showing morphology and scalp topography similar to the N2 component of the N2-P3 complex elicited by shifts of attention to infrequent salient events^12^ a reversal of the conditions historically subtracted to generate a difference wave (i.e., win-loss rather than loss-win) provided an emphasis on the relative positivity elicited by the win outcome (reviewed by ^7^). The RewP is thought to reflect receipt of midbrain dopaminergic reward prediction error signaling by the ACC, tracking magnitude and valence of expected vs. attained rewards and therefore encoding outcomes as better or worse than expected ^13^. As such, the RewP may subserve reinforcement learning ^6^ as an electrophysiological index of prediction error-based reward valuation ^5,8^. Although ERP studies of reward processing have largely focused on the RewP to date, interest is gaining in expanding assessments to include later components that reflect affective responses to reward outcomes^3^, such as the late positive potential (LPP)^3,14^, as well as components preceding the outcome that reflect reward anticipation, such as the stimulus-preceding negativity (SPN), a measure of anticipatory attention^3,15,16^. And more granular efforts to parse anticipatory from consummatory sub-stages of reward processing with ERPs^17^ have revealed ERPs that selectively covary with traits relevant for reward processing (e.g.,^18^).

In addition to ERPs, time-frequency measures of neuro-oscillatory power and phase synchrony have been useful in characterizing component processes related to reward. Frontomedial theta oscillations, more broadly associated with cognitive control and error processing ^6,19^, are responsive to anticipatory and evaluative phases of reward processing, particularly in association with losses and loss-related learning ^20-25^. In contrast, posterior delta oscillations are maximally responsive to winning outcomes^20,21^, possibly in connection with positive prediction error formation and subsequent behavioral adjustments ^26^. More specifically, prior work examining frequency-specific neuro-oscillatory EEG activity within the RewP time window has demonstrated that delta and theta evoked power are associated with ERPs to wins and losses, respectively ^27-29^ providing convergent evidence that neural mechanisms underlying feedback processing differ by outcome (but see^30^). The potential practical value of combining ERP and time-frequency measures is illustrated by a recent example from the clinical literature in which combining time frequency-based measures of delta oscillations with the RewP enhanced the sensitivity and positive predictive value of models for depression risk^31^.

Many previous neuroimaging studies examining reward processing have used paradigms that deliver rewards based on the participant’s behavioral performance and/or decision making such that rewards earned depend on participant action selection and motor responding (e.g. ^32^). Studies based on these and similar tasks make valuable contributions in modeling reward attainment under conditions requiring these higher-order processes. For example, meta-analysis indicates the RewP is positively modulated by the extent of personal control over outcome ^33^. Nevertheless, performance-based reward tasks pose an inherent challenge in isolating more basic reward processes from other features such as the individual’s motivation, cognitive effort, and performance skill. Modeling more basic reward features is not only relevant to understanding healthy reward functions, but is necessary to develop a more precise understanding of reward system dysfunctions in neuropsychiatric disorders with distinct profiles of reward-related deficits, or with co-occurring cognitive and motivational impairments ^34-36^.

It has previously been demonstrated that a RewP can be elicited by slot machine-style tasks in which reward outcomes do not depend on successful decision-making or motor responding ^37,38^, offering a compelling, real-word analog of elemental reward processes. Slot machine play is a common and highly reinforcing form of gambling that typically features near miss outcomes, in addition to wins and complete losses. Near misses are outcomes that have closer proximity to a win than a “total” loss, but have the same economic value as losses; they are typically experienced as more aversive but also more motivating than wins in that they increase play persistence ^39-41^. Given that near misses come closer to wins than other losses, the concept of frustrative non-reward, in which failure to obtain a desired goal invigorates behavior ^42^, has been invoked in explaining the near miss effect ^41^. Here, we used a slot machine paradigm to evaluate basic aspects of reward anticipation and outcome processing, including responses to near misses. The goals of the present study were to use a comprehensive set of EEG measures to parse psychologically distinct phases of reward processing. Accordingly, we combined analysis of ERPs reflecting anticipatory attention allocation (SPN) and early (RewP) and late (LPP) stages of reward outcome evaluation with time-frequency analysis (TFA) of theta and delta oscillatory total power and phase synchronization during time windows corresponding to the ERP measurements. After first characterizing the paradigm with time-domain ERP components, ERPs were then related to event-related magnitude changes (total power) and event-related phase synchronization and also to individual differences in age and anticipatory and consummatory reward sensitivity. In addition to establishing expected significant condition effects for the SPN, RewP, and LPP, we examined hypotheses that theta and delta measures would explain significant variance in the time-domain ERPs, and that this broad measurement set would explain variance in participant age and trait reward sensitivity.

## Methods

### Study Participants

Data were collected from 54 (42 male) healthy adults between the ages 19 and 61 (33.72 ± 14.4), recruited from the community through online advertisements, flyers, and word-of-mouth. The Structured Clinical Interview for DSM-IV (SCID-IV-TR) ^43^ ruled out current or past Axis I psychiatric disorders. Recent substance use for common testable drugs of abuse (e.g., cannabinoids, opiates, cocaine, amphetamines) was ruled out through urinalysis on assessment days. English fluency, and 18-65 years of age were required. History of head injury, neurological illness, or other major medical illnesses that affect the central nervous system were exclusionary criteria. All participants provided written informed consent under procedures approved by the Institutional Review Board at University of California, San Francisco (UCSF).

### Slot Machine Task

Participants completed 288 trials of a slot machine reward processing task developed for this study and informed by prior work ^37,38,44-46^. To build expectancy, the display consisted of 3 sequentially populated slot reel positions (stimulus onset asynchrony; SOA = 1.2 sec). Each reel was initially blank until populated with one of 12 possible fruit symbols, culled from a royalty free-clip art library, https://openclipart.org. Fruit symbols were distributed equally among possible outcomes, such that individual fruit symbols carried no predictive information about likelihood of winning. Total trial time after the participant initiated a given play was 6115 ms (see Figure 1 for detailed trial timing).

**Figure 1.**
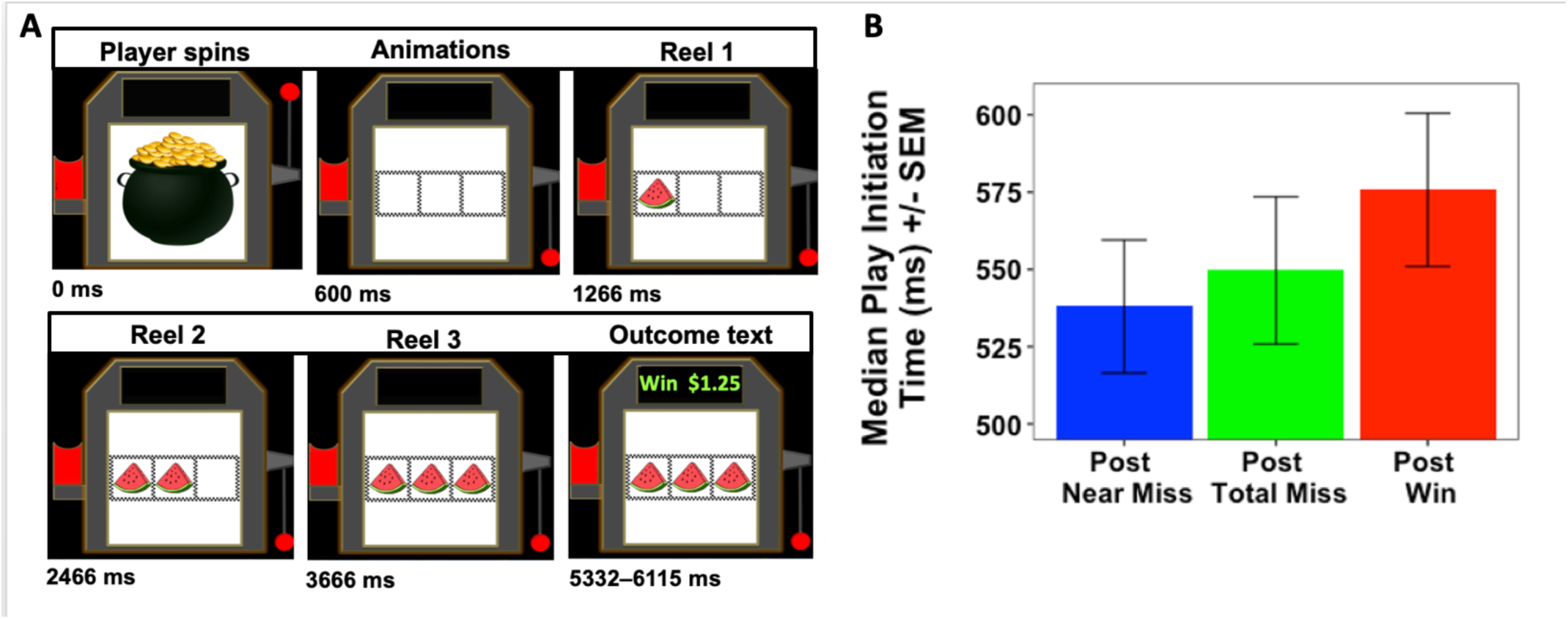
Slot machine task timing diagram: evolution of a single trial. Figure, Right shows a bar graph of player initiation times by trial type.

To increase face and ecological validity, specific task design features were included to mimic structural characteristics common to real-word slot machines, including sound effects (audible coin drop) and visualizations (coin insertion, lever pull, visual flicker during play outcome) ^40,47^. In order to promote a sense of agency and control which positively modulates the signals being studied^33,44^, trials were initiated via participant button press, after which timing of the slot reels was automated so that reward outcomes did not depend on participant decision-making or motor response preparation or execution. More specifically, the participant’s button press triggered an audible animated coin drop and lever press, followed by reel positions being populated sequentially, whereby reels 1 (R1), R2, and R3 were each populated with a single fruit symbol. After R3 populated, the outcome (reward evaluation) phase began and a visual checkerboard border flickered for 1000 ms followed by outcome text that depicted either “WIN $1.25” or “LOSE,” depending on the trial type. There were 3 trial types, presented in a pseudorandom sequence: frequent losses evident at R2 (total miss probability =.50), infrequent wins evident at R3 (win probability =.25), and infrequent near misses evident at R3 (near miss probability =.25). Wins (AAA) occurred when 3 identical fruit symbols were populated in the 3 slot reels; Near Misses (AAB) occurred when the first and second reel symbols matched but the R3 symbol did not match (AAB); Total Misses (ABC) occurred when the R2 symbol did not match the R1 symbol, indicating a loss at R2, with the R3 outcome providing no additional information about the loss (which is why Total Miss trials were time-locked to R2 for analysis). Near Miss and Total Miss trials were $0 payouts, whereas each Win trial yielded a $1.25 payout. To ensure that participants would feel that the opportunity for reward was valid, they played for a monetary bonus that reflected a portion of their slot machine winnings above their regular compensation for study participation.

### Self-Report of Hedonic Trait Sensitivity

The Temporal Experience of Pleasure Scale (TEPS) is self-report scale used to measure anticipatory and consummatory experiences of pleasure on a scale from 1 (“very false for me”) to 6 (“very true for me”) ^48^. Higher scores indicate higher levels of experiential pleasure (total score range 0-28). The anticipatory subscale (10 items; alpha = .76) primarily reflects receptiveness to rewards (e.g., “wanting”, and the consummatory subscale (8 items; alpha = .64) reflects hedonic response (e.g. “liking”) and inversely relates to anhedonia. Test-retest reliability over ∼6 weeks for the subscales was .80 and .75, respectively^48^.

### EEG Data Acquisition and Processing

EEG data were recorded from 64 channels using a BioSemi ActiveTwo system (www.biosemi.com). Electrodes placed at the outer canthi of both eyes, and above and below the right eye, were used to record vertical and horizontal electro-oculogram (EOG) data. EEG data were continuously digitized at 1024Hz and referenced offline to averaged mastoid electrodes before applying a 0.1Hz high-pass filter using ERPlab ^49^. Data were next subjected to Fully Automated Statistical Thresholding for EEG artifact Rejection (FASTER) using a freely distributed toolbox ^50^. The method employs multiple descriptive measures to search for statistical outliers (> ±3 SD from mean): (1) outlier channels were identified and replaced with interpolated values in continuous data, (2) outlier epochs were removed from participants’ single trial set, (3) spatial independent components analysis was applied to remaining trials, (5) outlier components were identified (including components that correlated with EOG activity), and data were back-projected without these components, and (6) outlier channels were removed and interpolated within an epoch. The original FASTER processing approach was modified between steps 2 and 3 to include canonical correlation analysis (CCA). CCA was used as a blind source separation technique to remove broadband or electromyographic noise from single trial EEG data, generating de-noised EEG epochs. This approach is identical to the CCA method described in our prior reports ^51,52^. Of the 288 trials presented, 278.19 +/- 10.93 were acceptable after artifact rejection and were subjected to analysis; condition-specific breakdown of trials subjected to analysis were: 69.67 +/- 3.14 wins, 69.78 +/- 3.02 near misses, and 138.74 +/- 5.50 total misses.

### ERP Measurement

Reel 3. Epochs were time-locked to the onset of R3 and baseline corrected using either the -100 to 0ms baseline preceding R3 for the RewP and LPP, which were measured post-R3 or the -2400 to -2300ms period preceding R3 (i.e., -100 to 0ms preceding R2) for the SPN, which was measured pre-R3. ERP averages were generated using a trimmed means approach, excluding the top and bottom 10% of single trial values at every data sample in the epoch before averaging to produce a more robust mean estimation ^53^. The SPN was measured as the average area in the window from -100 to 0ms prior to the R3 outcome ^15,16,37,38^, with a separate ERP average of the AA trials (collapsed across AAA and AAB trials) and the AB trials (i.e., the ABC trials). The RewP was measured as the average voltage between 228 – 344ms post R3 onset in the AAA-AAB difference wave, based on a time window established by meta-analysis of the RewP across 54 studies ^33^. The LPP was calculated as the average voltage between 600-800ms ^54^ after R3, and was measured for both wins and near misses separately (and compared statistically to ABC total misses).

Reel 2. In addition to the R3-locked RewP, a RewP was measured as the average voltage between 228 – 334ms post R2 in the AA-AB difference wave in order to compare the brain’s response to outcomes revealed at R3 (i.e., win AAA vs. near miss AAB) with outcomes revealed at R2 (i.e., possible win AA vs. total miss AB). LPP amplitudes were not similarly measured in response to R2 because the LPP time window overlaps with the SPN.

For statistical analyses, measurements were based on representative electrodes commonly used for specific ERP components in the literature (FCz for RewP ^12,14,37,38^, Cz for SPN^37,38^, and Pz for the LPP^3^).

### Time-Frequency Analysis (TFA)

Time frequency analysis of single trial EEG data was done with Morlet wavelet decomposition using FieldTrip software ^55^ in Matlab. Specifically, we used a Morlet wavelet with a Gaussian shape defined by a ratio (σ_f_ = f/C) and 6σ_t_ duration (f is the center frequency and σ_t_ = 1/(2πσ_f_). In this approach, as the frequency (f) increases, the

spectral bandwidth (6σ_f_) increases. Center frequencies were set to minimize spectral overlap for two frequency bins: delta = 3 Hz (range: 2.5-3.5 Hz) and theta = 5 Hz (range: 4.2-5.8 Hz). Inter-trial coherence (ITC) of phase was calculated as 1-minus the circular phase angle variance ^56^. ITC provides a measure of the phase consistency of frequency specific oscillations with respect to stimulus onset across trials on a millisecond basis. Total power was calculated by averaging the squared single trial magnitude values in each frequency bin on a millisecond basis. The average power values were 10log_10_ transformed and then baseline corrected by subtracting the mean of the pre-R2 stimulus baseline (−250 to -150 ms) from each time point separately for every frequency. The resulting values describe change in total power relative to baseline in decibels (dB). For each participant, time-frequency measures (power, ITC) were extracted from delta (3 Hz) and theta (5 Hz) bands in the same time windows corresponding to the ERP components of interest (i.e., SPN, RewP, LPP) and subjected to analyses as described below.

### Data Analysis Plan

#### Task behavior

For behavioral data, repeated measures analysis of variance (ANOVA) was used to compare median response time to initiate trials after wins (AAA), near misses (AAB) and total misses (ABC). Bonferroni correction was applied to control familywise type 1 error at p<.05 for pairwise follow-up comparisons across the three conditions.

#### ERP and TFA Condition effects

ERP amplitude measures for the SPN (AA vs. AB) and RewP (AAA vs. AAB) were assessed with t-tests, while the LPP was assessed with a repeated measures ANOVA (AAA vs ABC and AAB vs ABC). TFA difference score measures (theta power and ITC, delta power and ITC), within time windows corresponding to the RewP, were also assessed for condition effects with t-tests. Unlike the ERP waveforms, which exhibited expected complex morphologies that greatly differed between R2 and R3, the time-frequency decomposition produced simplified waveforms that enabled examination of single condition effects, to assess the impact of the three reward outcomes (win AAA at R3, near miss AAB at R3, total miss AB at R2) on underlying signals. Therefore, we conducted 4 single condition TFA ANOVAs (each with three levels of outcome). Total miss (AB) trials were downsampled to avoid confounding trial number in these comparisons. Multiple comparisons correction strategy: Our evaluation of ERP and TFA condition effects yielded 7 EEG measure comparisons with which to assess the primary condition effects of interest (SPN, RewP, repeated measures LPP (wins, near misses), and repeated measures theta and delta power and ITC (wins, near misses, total misses). Bonferonni-correction was applied to account for these 7 comparisons to control for type 1 error across the measurement set, with each model held to a threshold of p < .0071.

#### Correlation analyses

##### ERP and TFA Relationships

The relationships between ERP and TFA difference score measures were assessed using separate parametric multiple regression models predicting each of the four ERP measures (SPN, RewP, LPP_win, LPP_nm) with the four delta and theta TFA measures from the corresponding ERP time window.

##### Participant Age

Given our relatively wide age range (19-61 years) we assessed the association of age with the electrophysiological measures of reward processing using bonferonni-corrected Pearson product-moment correlation coefficients.

##### Reward sensitivity

Relationships between electrophysiological and self-report measures of reward sensitivity (TEPS anticipatory and TEPS consummatory) were assessed with multiple regression models.

## Results

### Task Behavior

Response times: There was a significant difference among response times to initiate trials following a win (AAA), near miss (AAB), or total miss (ABC) outcome (condition effect omnibus p = .003) (Table 1). Follow-up comparisons among the conditions indicated that participants were faster to initiate trials following a near miss, relative to a win (bonferroni-corrected, p=.003) while response times did not significantly differ among any other pairwise comparisons.

**Table 1.**
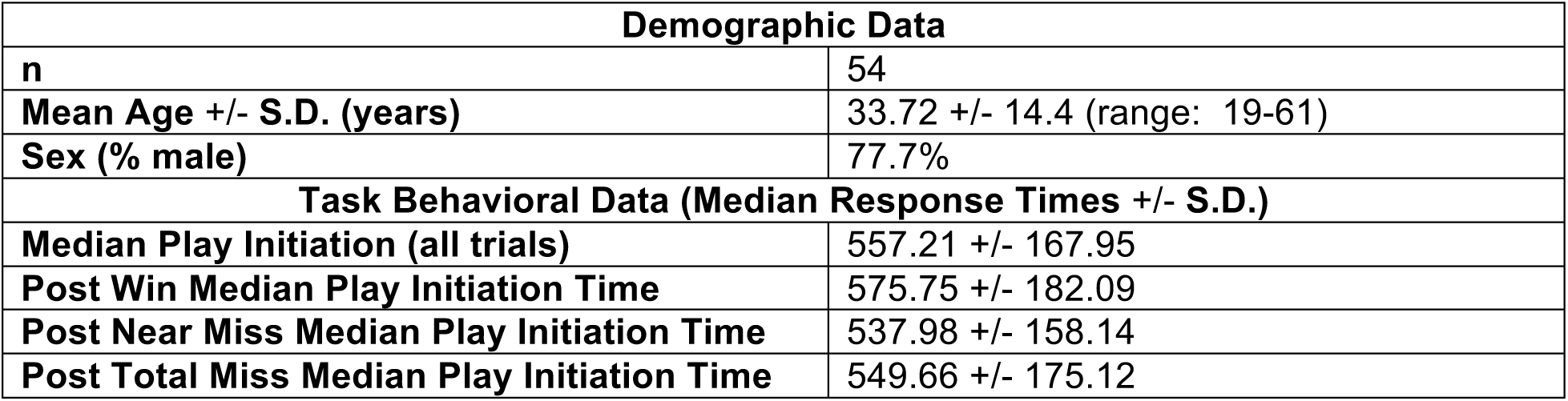
Participant demographic characteristics and play initiation times.

| Demographic Data |  |
| --- | --- |
| n | 54 |
| Mean Age +/- S.D. (years) | 33.72 +/- 14.4 (range: 19-61) |
| Sex (% male) | 77.7% |
| Task Behavioral Data (Median Response Times +/- S.D.) |  |
| Median Play Initiation (all trials) | 557.21 +/- 167.95 |
| Post Win Median Play Initiation Time | 575.75 +/- 182.09 |
| Post Near Miss Median Play Initiation Time | 537.98 +/- 158.14 |
| Post Total Miss Median Play Initiation Time | 549.66 +/- 175.12 |

### ERP Effects

Anticipation (SPN): Prior to R3 onset, a characteristic SPN with a central topography was observed for potentially winning trials (AA; i.e., trials not yet revealed as a win or near miss), relative to total miss (AB) trials (Figure 2). There was a significant effect of condition, driven by more negative SPN amplitudes at Cz when comparing AA to AB trials (t (53) = -5.22, p<.001).

**Figure 2.**
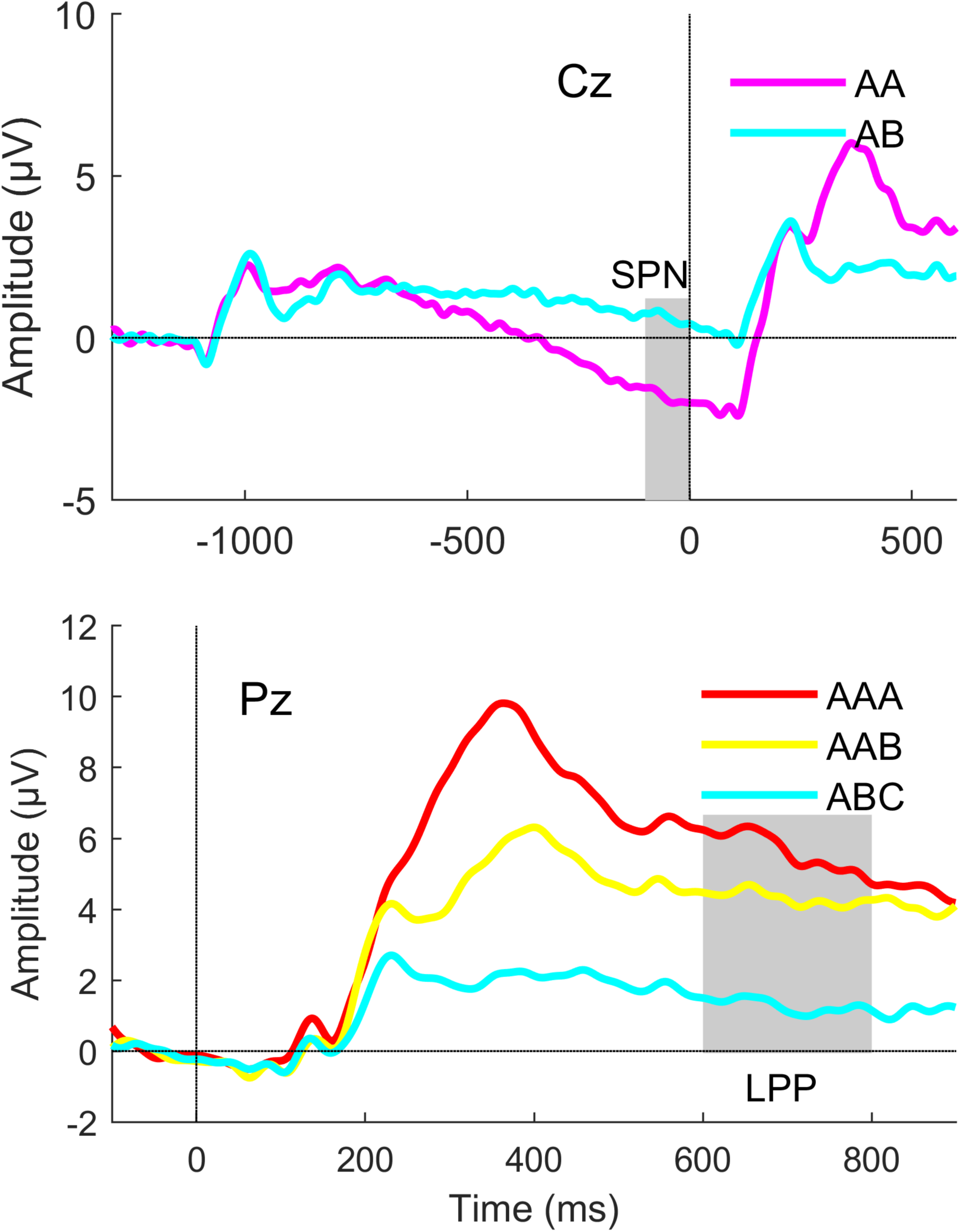
Top: Stimulus preceding negativity (SPN) grand average waveforms at electrode Cz, depicting Reel 2 time-locked possible win (AA) trials, shown in pink and total miss (AB) trials, show in cyan. Bottom: Late positive potential (LPP) grand average waveforms at electrode Pz for win (AAA) trials shown in red and near miss (AAB) trials shown in yellow. Time 0 ms corresponds to Reel-3 outcome. Grey bars depict ERP measurement window; n = 54.

Early Outcome Evaluation (RewP): Grand average waveforms (Figures 2 and 3) show a RewP with expected frontocentral topography in the difference wave of wins (AAA) – near misses (AAB) peaking in the measurement window (228 – 334ms) ^33^. There was a significant difference at FCz, with ERP amplitude to wins being larger than to near misses (t(53) = 8.76, p<.001). In addition, the RewP difference score was larger at Reel 3 (AAA-AAB) than Reel 2 (AA-AB) t (53)=8.40, p<.001).

**Figure 3.**
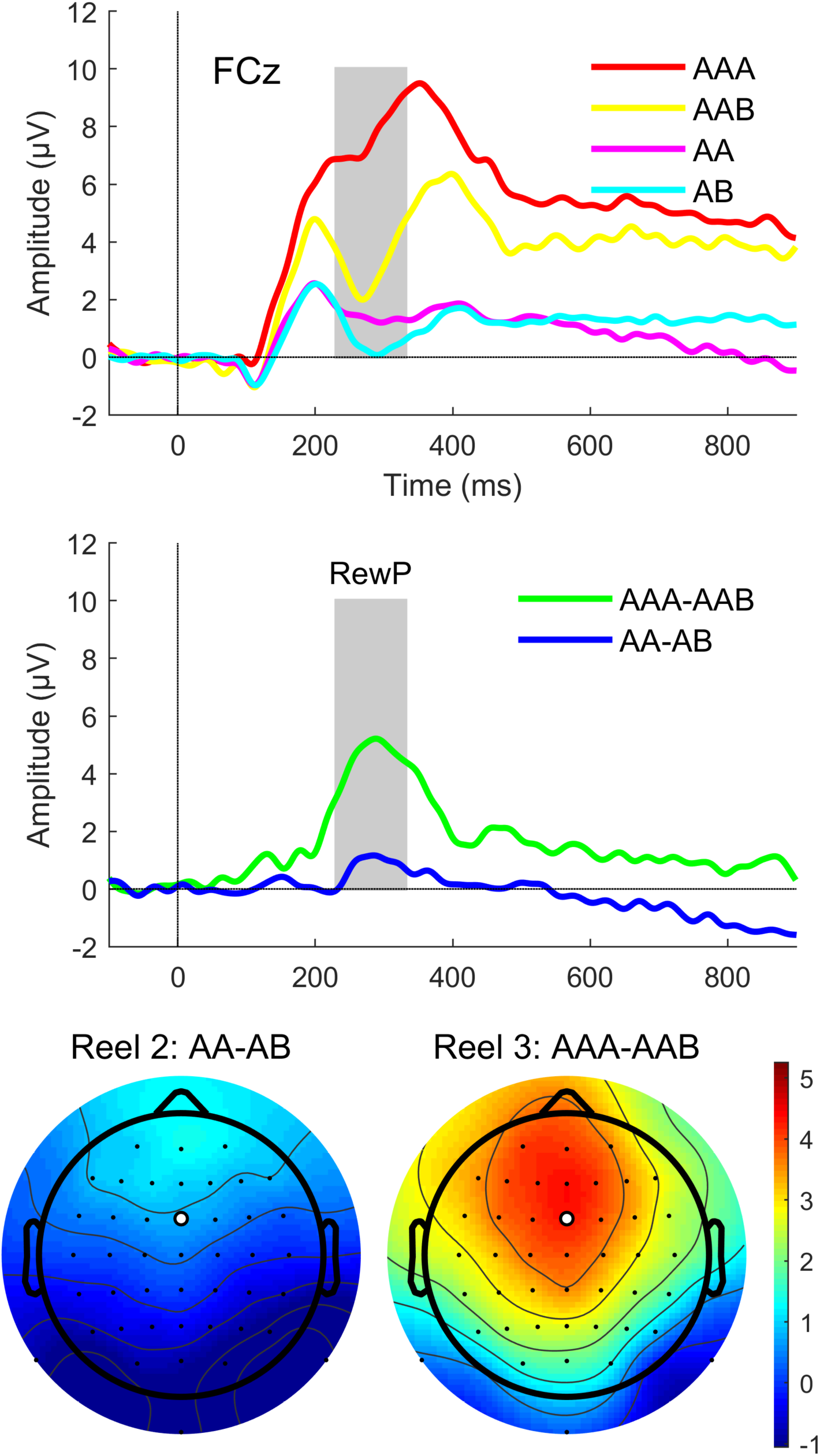
Effects of Reel Position. Top: Grand average ERP waveforms by condition at electrode FCz. Waveforms for Reel-3 locked (AAA, AAB) and Reel-2 locked (AA, AB) individual conditions. 0 ms on the x-axis corresponds to Reel-2 (AA, AB) outcome, which spanned from 1494-2000 ms from trial onset and Reel-3 (AAA, AAB) outcome which spanned from 3894-4000 ms from trial onset. Middle: Difference waves for Reel 3-locked AAA-AAB and Reel 2-locked AA-AB comparisons. Grey bar depicts ERP measurement window for Reward Positivity (RewP). Bottom: Topographical maps of condition difference waves at Reel 2 (bottom, left) and Reel 3 (bottom, right).

**Figure 4.**
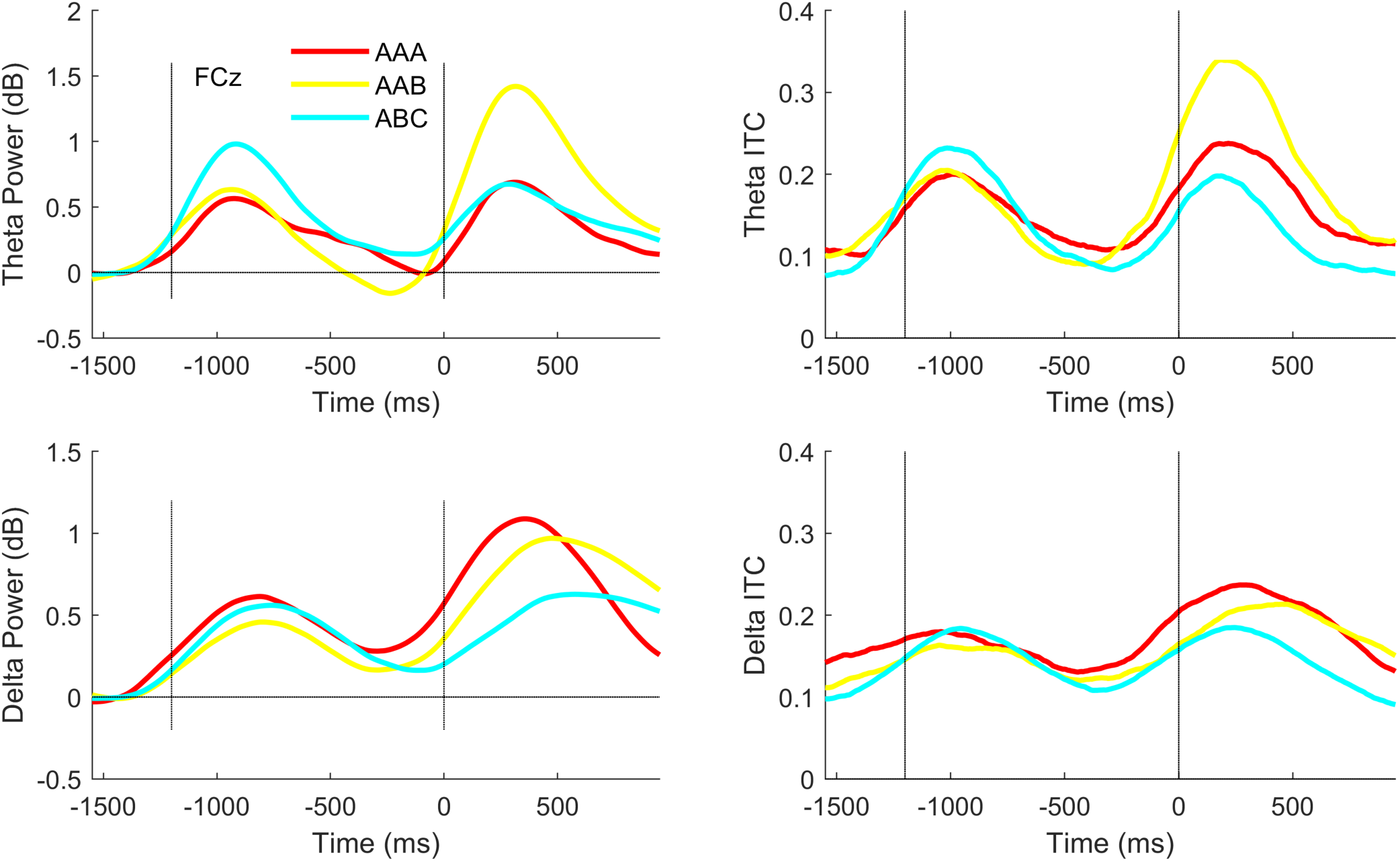
Condition effects for total power (left) and inter-trial coherence (ITC) (right), for theta (5Hz; Top) and delta (3Hz; Bottom) bands. Time 0 ms corresponds to Reel-3 outcome. ABC trials were downsampled to equate trial numbers across conditions.

Later Outcome Evaluation (LPP): Grand average waveforms (Figure 2) show a sustained positivity with a posterior distribution. LPP showed a significant main effect of condition at Pz: F(2, 106) = 58.12, p<.001). Planned single degree of freedom follow-up contrasts indicated LPP_wins (AAA) differed significantly from LPP_total misses (ABC), p <.001 and similarly LPP_near misses (AAB) differed significantly from LPP_total misses (ABC), p = .001.

### Time-Frequency Analysis

We observed significant condition effects for both theta power and ITC (omnibus p<.001 for both theta power and ITC). Bonferroni-corrected follow-up tests across the three outcomes indicated differences were explained by increased power and phase synchrony in response to near misses, relative to both wins (power t(53)=4.06, adjusted-p=.001; ITC t(53)= 5.17, adjusted-p<.001) and R2 total misses (power t(53)=2.99, adjusted-p=.033; ITC t(53)= 5.48, adjusted-p<.001), and that theta activity did not distinguish R3 wins from R2 total misses. Delta power and ITC also showed significant condition effects (omnibus delta power p = .007; delta ITC p = .003), with follow-ups indicating greater delta power and ITC to R3-wins relative to R2-total misses (delta power t(53)= 3.03, adjusted-p = .01; delta ITC t(53)= 3.42, adjusted-p = .003). R3-near misses showed numerically intermediate values of delta power and ITC, falling between R2-total misses and R3-wins but not significantly differing from either (adjusted-p> .12). See Figure 3.

### Time-Frequency Difference Score Measures as Predictors of ERPs

Regression models of ERP component difference scores (SPN, RewP, LPP_wins, LPP_near misses) on theta and delta signals within time windows corresponding to each ERP component were conducted in order to determine the underlying contributions of delta and theta total power and phase synchrony to the time-domain ERP component effects associated with different stages of reward processing. The overall model regressing RewP difference scores on theta power, delta power, theta ITC and delta ITC difference scores (all measured in the RewP time window at FCz) was significant F(4, 49)= 5.55, p=.001; R^2^=.31. Theta ITC (t=-2.63, p=.01) and delta ITC (t= 2.05, p=.046) difference scores both made significant unique contributions in the model, with delta phase synchrony difference scores showing a positive relationship with the RewP, and theta phase synchrony difference scores showing a negative relationship. Neither power measure difference score was a significant regressor in the model (p>.05).

In contrast, the regression models predicting SPN or LPP amplitude from TFA measures within corresponding time windows was not significant (SPN: F (4, 49)= 1.34, p=.27; R^2^= .10; LPP to AAA: F (4, 49)= 0.61, p=.66; R^2^= .05; LPP to AAB: F (4, 49)= 1.08, p=.38; R^2^= .08), indicating these theta and delta time-frequency measures were not a significant predictor of SPN or LPP ERPs.

### Relationships with Age

Age relationships were examined among the four ERP (SPN, RewP, LPP_wins, LPP_near misses) and theta and delta power and ITC to wins, near misses, and total misses) using a bonferonni-corrected alpha across these measures (p<.003). There was a significant negative association between the RewP and age (r=-0.46; p =.001); no other comparisons surpassed the adjusted statistical significance threshold (.03<|r|<.25 ; 0.07<p<.82).

### Relationships with Behavioral Measures of Reward Sensitivity

Relationships between electrophysiological and self-report measures of reward sensitivity were assessed in 2 regression models (TEPS anticipatory and consummatory pleasure), with bonferonni-correction applied, across the two models (p<.025). The first model regressed the TEPS-anticipatory onto the ERP anticipatory measure, the SPN, which was not a significant regressor (t= 0.98, p=.93; model R^2^ = .01). The second model regressed the TEPS-consummatory measures onto the remaining ERP measures, which represented post-outcome responses (RewP, LPP to wins, LPP to near misses). Here, the LPP_win was positively associated with variance in TEPS consummatory scores (t= 2.79, p=.007; model R^2^= .14), while betas for the RewP (t= -.57, p=.57) and LPP_near miss ((t= -2.02, p=.05) were not significant.

## Discussion

The goal of this study was to extend prior research on reward-related brain functioning by extracting EEG measures of reward anticipation and early and late-stage reward outcome evaluation in the same paradigm, combining ERP and TFA measures in characterizing these sub-components of reward processing, and relating these neurophysiological responses to psychologically distinct phases reflecting wanting (anticipation) vs. liking (consummation). After replicating expected condition effects for ERP and TFA measures of interest, key findings of this study i) extend the literature by demonstrating that delta and theta phase synchrony measures are correlated with the RewP, but not SPN or LPP, amplitude, ii) show that the measures under study covary selectively with participant age and self-reported sensitivity to consummatory reward processes, iii) demonstrate that the RewP is larger at R3 than R2, and iv) show that near miss outcomes induce larger and more synchronized frontomedial theta responses than wins and total misses. The slot machine context, which minimizes cognitive and motivational demands, may be particularly relevant for assessing basic reward responsivity in clinical or developmental populations in which within-group variation or case-control differences in motivation and cognition can complicate interpretation of performance-based reward tasks.

Consistent with the literature, we found an SPN precedes R3 on possible win (AA) but not definite loss (AB) trials. This was first modeled by Donkers in a 3-reel slot paradigm, demonstrating that an SPN precedes, but does not follow, outcome feedback ^37,38^, and supports the interpretation that, in the context of a reward paradigm, the SPN reflects reward anticipation ^15,16^. The SPN comprises a class of slow negative potentials thought to signify a generalized attentional control response generated during anticipation of impending feedback across varied experimental paradigms ^15,16^. While the SPN is not exclusively generated by incentivized contexts, it is strongly influenced by motivational content of the anticipated feedback ^57,58^.

The presence of a RewP with characteristic topography and time signature (Figure 3) is strongly expected by prior literature across varied reward task paradigms. These are reviewed in ^5-8^ and include studies of slot machine play most relevant to the current study ^37,38,59-61^. The RewP was also the only measure examined that significantly covaried with age, with younger adults showing larger amplitudes. Many prior studies examining covariation with the RewP (or separate win and loss waveforms) have focused on developmental samples, though studies that have extended the age range into middle and older adults observe a pattern similar to ours, with attenuation of the RewP over the course of adulthood^62,63^. Several existing studies used a slot paradigm design with all outcome information being delivered simultaneously (with trial outcomes defined based on number or proximity of stimuli in the final visual array) ^59-61^. In contrast, our task design utilized sequential reels enabling us to examine the effects of reel position on outcome processing, when feedback about loss is revealed sequentially over time, ostensibly allowing anticipation for possible win trials to further develop. Prior electrophysiological studies have shown that ERP components relevant for reward evaluation such as the RewP are modulated by reward proximity, suggesting that “close” outcomes such as the near miss may be processed distinctly from both win and “total miss” events ^59,60,64,65^. Near misses have also been associated with theta oscillations in reward network regions including the insula and right orbitofrontal cortex ^45^. However, the R3-locked RewP contains both the reward-related positivity to the win as well as the relative negativity to the near miss. Thus, both may contribute meaningful variance to the RewP as a difference score^66^. Because our task design enabled comparing temporally distinct reel outcome effects in the RewP time window, we were able to observe that the RewP is significantly larger at R3 (AAA-AAB) than R2 (AA-AB), despite the equivalent conditional probability of losing at each of these reels (and the equivalent economic valuation of a zero payout). Observing a R2 RewP converges with prior work demonstrating the RewP can be elicited by intermediate feedback using a performance-dependent (modified monetary incentive delay) reward task ^67^. However, though the slot task in the current study produces two RewP components, the R3-locked one is significantly larger suggesting that the win on R3 drives the R3 RewP amplitude increase more than the losses (which are present at both reels), fitting with the conceptualization that the RewP predominantly represents win-related signaling ^7,12^.

We also observed an LPP after the RewP that was similarly affected by trial outcome, being larger to wins than to near misses. The LPP has most often been studied in the context of affective processing (e.g., affective picture exposure) and is thought to signify attentional allocation to stimuli with affective content, possibly in the service of facilitating encoding of emotionally relevant stimuli ^54^. LPP amplitude covaries with BOLD activations in temporo-parietal and occipital cortices ^68^ consistent with the component’s characteristic posterior scalp topography. However, studies using temporal-spatial PCA have indicated that the large, slow positivity characterized by the LPP can be fractionated into sub-components that have distinct spatial morphologies (e.g., occipital vs. parietal) and may respond selectively to cognitive vs. affective aspects of task stimuli ^28,69,70^. This distinction is supported by Cockburn et al, who focused on the role of feedback quality on RewP amplitude elicited during a time estimation task and found that a posterior slow wave (500-700ms, post-feedback) became more positive as feedback became more informative ^14^. The authors suggested the slow, late wave they observed could signify higher-order encoding of feedback that may functionally apply the information carried by earlier evaluative components like the RewP to formal learning principles as outcomes are further processed. Indeed, this interpretation is consistent with the idea that the RewP, as an initial reward evaluation response, signals receipt of the reward prediction error into ACC, while later activities may be more tightly coupled to subsequent behavioral adjustment based on the initial processing of feedback ^71^. Though our study had no manipulation of feedback quality or other cognitive manipulation, the LPP to wins significantly related to self-reported consummatory reward behavior. In our data, the LPP response to wins was strongly related to the SPN to possible wins (p<.001; data not shown) with larger LPP to wins co-varying with more negative SPN amplitudes, suggesting that, similar to a prior report using a dissimilar reward task ^72^, SPN and LPP responses, though psychologically and temporally distinct, may share underlying functional features. Similarly, the SPN has been shown to co-vary with the feedback P3 in a monetary incentive delay task. Though counterintuitive, it is possible that the reason we do not observe an association between the SPN and anticipatory pleasure, is that the SPN covaries more with consummatory reward behavior.

Time-frequency analyses were undertaken on single trial EEG activity to examine the contribution of delta and theta oscillatory bands to observed ERP difference-score condition effects during anticipatory and feedback phases of slot play. First, we found that neither theta nor delta power or phase synchrony were associated with SPN or LPP amplitudes. In contrast, theta and delta phase synchrony measures were both significantly associated with RewP amplitude, with a stronger relationship for theta than delta. Thus, our data suggests that the RewP derives from phase resetting of theta and delta oscillations rather than a change in the magnitude of these oscillations. Our findings support prior conclusions based on analyses of evoked power^27-29^ and ITC ^27^ indicating that theta and delta make independent contributions to the time domain RewP. The single trial time-frequency analysis we undertook extends prior work based on time-frequency decomposition of trial averages by examining oscillatory total power and phase synchrony which offers a different account EEG dynamics ^73^ (e.g., by modeling spectral amplitude perturbations, regardless of stimulus phase). The RewP, or more specifically the FRN, has been theoretically and empirically tied to the more widely studied error-related negativity (ERN) ^74,75^ a frontomedial negativity elicited by internal error or conflict detection (as opposed to the external feedback that elicits a RewP/FRN); thus the ERN and RewP may share an underlying mechanism that functions as an error detection system ^11^. Though the RewP is typically defined as a difference score of wins vs. losses, resulting in a relative positivity (reviewed by ^7^, its amplitude has contributions from both the positivity to wins and the negativity to losses ^12^. Our data provide further evidence that there are unique contributions to RewP from theta and delta respectively

Additional time-frequency analyses isolated condition specific effects that would be difficult to disentangle in the temporal domain due to component overlap and large magnitude differences between R2 and R3. Condition-specific phase synchronization and magnitude changes in response to event onsets (ITC and total power) were observed, corresponding with increased delta oscillation magnitude to wins and increased theta oscillation magnitude and synchronization to near misses. Our findings support prior studies of reward outcome valence that associate theta with loss-related processing and delta with winning outcomes ^20,21,23,25,27-29^. Notably, near misses in our data elicited the largest magnitude and phase synchrony of the frontomedial theta response, with significantly more theta power and synchrony than wins as expected, but also significantly more than for total misses (i.e., losses revealed at R2). Furthermore, subsequent play behavior was influenced by outcome on the preceding trial as play initiation times following a near miss were significantly faster than initiation times subsequent to a win. This effect could be interpreted as a post-reinforcement pause following winning trials that has been described in the literature^76,77^. However, this effect is also consistent with the possibility of a frustration effect^42^ being induced by near misses that invigorates future play, as others have suggested in the context of gambling near misses ^41,77^. That our data demonstrate that R3 “near miss” loss elicits more theta signaling than total misses despite the equivalent economic valuation and conditional probability of both loss outcomes, highlights the sensitivity of theta signals to reward proximity, and suggests a possible electrophysiological mechanism underlying frustrative non-reward.

One study limitation is that theta and delta bands were chosen *a priori* based on prior evidence of involvement in reward-related processing ^20-23,26^; and so we cannot rule out the possibility that frequency bands that we did not examine explain variance in the SPN or LPP, or additional variance in the RewP. Future research in the frequency domain focusing comprehensively on all bands and time points, including cross-frequency coupling would be useful. In addition, combined EEG-fMRI analyses will be useful to characterize neuroanatomy relevant to these EEG measures. Taken together, these results highlight the differential contributions of ERP and TFA measures to psychologically distinct aspects of reward processes involved in anticipating vs. experiencing pleasure, and add to the literature characterizing reward processing features during slot play, including the near miss response.

## Acknowledgements

VA: CX001028 (SLF). Drs. Fryer, Ford, and Mathalon are employees of the U.S. Government. The content is solely the responsibility of the authors and does not necessarily represent the views of the Department of Veterans Affairs.

